# Astrocyte-mediated transduction of muscle fiber contractions synchronizes hippocampal neuronal network development

**DOI:** 10.1101/2022.04.12.487995

**Authors:** Ki Yun Lee, Justin S. Rhodes, M. Taher A. Saif

## Abstract

Exercise supports brain health in part through enhancing hippocampal function. The leading hypothesis is that muscles release factors when they contract (e.g., lactate, myokines, growth factors) that enter circulation and reach the brain where they enhance plasticity (e.g., increase neurogenesis and synaptogenesis). However, it remains unknown how the muscle signals are transduced by the hippocampal cells to modulate network activity and synaptic development. Thus, we established an *in vitro* model in which the media from contracting primary muscle cells (CM) is applied to developing primary hippocampal cell cultures on a microelectrode array. We found that the hippocampal neuronal network matures more rapidly (as indicated by synapse development and synchronous neuronal activity) when exposed to CM than regular media (RM). This was accompanied by a 1.4-fold and 4.4-fold increase in the proliferation of neurons and astrocytes, respectively. Further, experiments established that the astrocytes release factors that inhibit neuronal excitability and facilitate network development. Results provide new insight into how exercise may support hippocampal function through regulating astrocyte proliferation and subsequent taming of neuronal activity into an integrated network.

**Highlights:**

- Contracting muscle conditioned media enhances neuronal activity.
- Contracting muscle conditioned media expedites neuronal maturation and accumulation of filamentous actin at presynaptic terminals.
- Contracting muscle conditioned media induces significant neuron and astrocyte proliferation.
- Astrocytes release factors that inhibit muscle media-induced neuronal activity.

## Introduction

Exercise is a highly effective strategy for maintaining cognitive health throughout life, even when initiated at late stages in life (Churchill et al., 2002; Erickson and Kramer, 2009; Erickson et al., 2011). Many studies have shown robust long-term changes in the hippocampus from increased physical activity, such as increased adult hippocampal neurogenesis, synaptogenesis and enlarged hippocampal volume which likely support enhanced cognition (van Praag et al., 1999; Redila and Christie, 2006; Clark et al., 2009, 2011; Erickson et al., 2011). However, the mechanisms by which exercise produces such dramatic changes in the hippocampus remain elusive. Uncovering the mechanisms that are responsible for enlarging the hippocampus and enhancing its function could be used to reverse-engineer treatments for cognitive pathologies that result in a diminished size and function of the hippocampus, such as Alzheimer’s disease, stress, depression, anxiety, PTSD, Cushing’s disease, epilepsy, and normal aging (Dhikav and Anand, 2007).

Cumulative research over the past few decades has suggested that factors released from contracting muscles (such as lactate (el Hayek et al., 2019), growth factors (Trejo et al., 2001; Fabel et al., 2003), trophic factors (Church et al., 2016), and myokines (Wrann et al., 2013; Moon et al., 2016)) provide crucial signals that support enhanced plasticity (Delezie and Handschin, 2018). However, how muscle factors affect hippocampal cells is still being worked out. Recently, we found that repeated electrical contractions of the hindlimb muscles of anesthetized mice in a pattern that produced endurance adaptations in the muscles (40 reps, twice a week for 8 weeks) caused increased numbers of new astrocytes in the hippocampus and enlarged the volume of the dentate gyrus by approximately 10% (Gardner et al., 2020). This suggests astrocytes are sensitive to muscle factors and proliferate when they detect muscle factors in the blood. Given the role that astrocytes play in forming the blood-brain barrier, they are well situated to transduce signals from the blood into the brain.

One way to study the interactions between contracting muscle cells and hippocampal cells including neurons and astrocytes is to isolate the cells and perform experiments *in vitro*. For example, previous *in vitro* studies found that muscle conditioned media attracted neurites of spinal cord motor neurons to form neuro-muscular junctions (McCaig, 1986). Along this line, our lab has been examining cross-talk between muscles and neurons *in vitro.* We recently found that when media from contracting muscle fibers derived from a C2C12 mouse myoblast cell line is applied to neuronal cultures derived from a mouse embryonic stem cell line plated on a micro-electrode array, it enhanced overall neural firing rates of the neurons (Aydin et al., 2020).

To further explore how factors from contracting muscles might influence hippocampal cells, we developed an *in vitro* preparation in which primary mouse skeletal muscle cells are plated on a functionalized substrate. The myoblasts develop bundles of myotubes and begin to contract spontaneously. We then take the media surrounding the contracting muscles (conditioned media, CM) and apply that media to *in vitro* primary hippocampal cell cultures that include neurons and astrocytes. The objectives of this study were to determine whether CM influences the function and maturation of hippocampal neuronal networks, and to investigate the role of astrocytes in the process of transduction of muscle contractions to the activity of hippocampal neuronal networks *in vitro*.

## Experimental procedures

### Primary mouse skeletal muscle and hippocampus dissection

Muscle tissues from the hindlimbs of 4-week-old CD1 mice were collected and dissociated using a standard protocol (Wang et al., 2017) with slight modifications. Briefly, the tissues were collected in cold PBS (Corning), minced, and digested for 30 minutes in digestion media consisting of DMEM, 2.5% HEPES, 1% GlutaMAX (all from Gibco), and 1% Penicillin-Streptomycin (Lonza) with the addition of 400 unit/ml collagenase (Worthington) and 2.4 unit/ml dispase (Sigma). The remaining tissues were triturated by pipetting in 0.25% trypsin (Gibco), then filtered by 70 and 40 μm cell strainers. After the dissociation, the pre-plating technique (Rando and Blau, 1994; Qu et al., 1998) was implemented to remove fibroblasts and increase the yield of myoblasts. The pre-plating technique steps are following. First, the dissociated cells were plated and incubated in uncoated flasks for three hours. Second, the supernatant with floating cells was collected and transferred into functionalized culture dishes.

For hippocampal dissection, hippocampal tissues were isolated from 2-day-old CD1 mouse pups and dissociated into single cells by following established protocol (Seibenhener and Wooten, 2012). The dissected hippocampal tissues were collected in cold Hibernate-E (Gibco), minced, then digested twice in 2 mg/ml papain (Sigma) for 30 minutes. The remaining tissues were mechanically dissociated further by pipetting, then filtered by 70 and 40 μm cell strainers.

All procedures were approved by the University of Illinois Institutional Animal Care and Use Committee and adhered to NIH guidelines (protocol number: 21053).

### Cell culture

Culture dishes were coated with 0.1 mg/ml Matrigel (Corning) for preparations of cell culture. The primary skeletal myoblasts were maintained below 70~80% confluency in muscle growth media consisting of Ham's F-10 Nutrient Mix, 20% fetal bovine serum, 1% GlutaMAX, 1% MEM Non-Essential Amino Acids (all from Gibco), 1% Penicillin-Streptomycin, and 0.5% chick embryo extract (US Biological) with ice-cold 10 ng/ml bFGF, 20 μM forskolin, and 100 μM IBMX (all reagent from Sigma). Once the confluency reached 70~80%, then the culture was maintained in muscle differentiation media consisting of DMEM and Ham's F-12 Nutrient Mix at a volume ratio of 1:1, 10% horse serum, 1% GlutaMAX (all from Gibco), and 1% Penicillin-Streptomycin to initiate myotube formations. Once myotubes were matured and contractions were observed, the media was changed with pre-muscle conditioned media consisting of Advanced DMEM/F-12, 1% GlutaMAX (all from Gibco), and 1% Penicillin-Streptomycin.

For the hippocampal neuron culture, the preparation of cell culture is the same as muscle culture. Hippocampal neuron cells were cultured using specially formulated media by the lab such as RM, CM, astrocyte media conditioned by RM (Ast-RM), and astrocyte media conditioned by CM (Ast-CM) depending on experiments (see details in Materials and methods).

### Muscle anti-actomyosin and glia anti-mitotic treatments

To inhibit skeletal muscle contraction, the specific reagent was used as shown in previous studies (Cheung et al., 2002; Pinniger et al., 2005). In the fully differentiated muscle culture with spontaneous contraction, 10 μM *N*-benzyl-*p*-toluene sulphonamide (BTS) (Sigma) was added. After 5-6 days, the BTS solution was replenished with the control media with 0 μM BTS and tension recovery was monitored after the washout.

In specific experiments below where indicated, the proliferation of glia was inhibited using a standard protocol (Liu et al., 2013). Briefly, the culture was treated with a cocktail consisting of 20 μM 5-fluorodeoxyuridine (MP Biomedicals), 20 μM uridine, and 0.5 μM Arabinofuranosyl Cytidine (all from Sigma) on day 1 and incubated for 72 hours. After 72 hours, 2/3 of the media was replenished with new media without the cocktail. A day after, the whole media was changed without the cocktail.

### Collection of muscle and astrocyte conditioned media

For the collection of RM and CM, primary skeletal myoblasts were cultured in muscle growth media, then muscle differentiation media when the confluency reaches 70~80%. Once myotubes were matured and 10~20% of myotubes began to twitch autonomously which is about 4 days after, the culture was maintained in pre-muscle conditioned media. The pre-muscle conditioned media was collected using 0.22 μm filters every 24 hours for 8 days and stored at −80 °C. For control, pre-regular media was collected from a culture dish without muscle cells, treated, incubated, and stored the same way as the CM. The final forms of RM and CM consist of pre-regular and muscle conditioned media, respectively and Neurobasal medium at a volume ratio of 1:1, 10% KnockOut serum replacement, 1% GlutaMAX (all from Gibco), and 1% Penicillin-Streptomycin with ice-cold 0.1◻mM β-mercaptoethanol (Gibco), 10◻ng/ml glial-derived neurotrophic factor (Neuromics), and 10◻ng/ml ciliary neurotrophic factor (Sigma).

For the collection of Ast-RM and Ast-CM, primary hippocampal neurons and astrocytes were cultured in basal media consisting of Advanced DMEM/F-12 and Neurobasal medium at a volume ratio of 1:1 10% KnockOut serum replacement, 1% GlutaMAX, and 1% Penicillin-Streptomycin. Once the confluency reached 100%, the media was replenished with RM and CM for the collection of Ast-RM and Ast-CM, respectively. The media was collected using 0.22 μm filters every 24 hours and stored at −80 °C.

### Immunofluorescence

Immunocytochemistry was performed by following steps. Samples were fixed with 4% paraformaldehyde, permeabilized with 0.05% Triton-X for 15 minutes at room temperature, and blocked with buffer solution consisting of PBS, 5% goat serum (Sigma), and 1% bovine serum albumin (Sigma) overnight at 4 °C. The samples were treated with primary antibodies overnight at 4 °C, secondary antibodies for 2 hours, and DAPI (1:1000; Invitrogen, D1306) for 20 minutes at room temperature.

Primary antibodies were anti-synaptophysin monoclonal rabbit (1:1000; Abcam, ab32127), anti-PSD95 monoclonal mouse (1:1000; Invitrogen, MA1-046), anti-Bassoon monoclonal mouse (1:1000; Abcam, ab82958), anti-β III Tubulin polyclonal rabbit (1:1000; SYSY, 302 302), anti-NeuN monoclonal mouse (1:1000; Abcam, ab104224), anti-Nestin polyclonal chicken (1:1000; Abcam, ab134017), anti-S100β monoclonal mouse (1:1000; SYSY, 287 011), anti-S100β polyclonal chicken (1:1000; SYSY, 287 006), and Alexa Fluor 647-conjugated Phalloidin (1:500; Invitrogen, A22287). Secondary antibodies were goat anti-chicken IgY Alexa Fluor 488 (ab150173), goat anti-mouse IgY Alexa Fluor 488 (ab150117), goat anti-rabbit IgY Alexa Fluor 568 (ab175696), and goat anti-mouse IgY Alexa Fluor 647 (ab150119) (all 1:500; Abcam).

All samples were imaged using Zeiss 710 confocal microscope (Carl Zeiss Microscopy).

### Synapse detection using double-fluorescent label method

To detect and measure synapses and filamentous actin at presynaptic terminals, the double-fluorescent label method was used by following protocol with modifications (Dzyubenko et al., 2016). Briefly, synapses or F-actin were double-labeled with two different antibodies at different channels. One set of puncta from one channel and the other set from the other channel were colocalized, then verified as synapses. The total intensity of synapse and F-actin was measured from integrated image planes after the colocalization process. The analysis was performed by ImageJ.

### Calcium Imaging

Calcium imaging was performed using Cal-590-AM (AAT Bioquest, 20510) by the manufacturer’s protocols. Briefly, samples were incubated with DMEM, 0.04% Pluronic F-127 (Sigma), and 5 μM Cal-590-AM for an hour at 37°C. After washout, samples were supplemented with DMEM with no phenol red to reduce background noise. The dye-loaded cells were excited at 574 nm and imaging was performed at the frame rate of 12 fps. For quantitative analysis, the average fluorescence intensity of selected regions of interest was calculated. Then the trace of fluorescent dynamics was calculated as Δ*F*/*F*_0_ = (*F_n_* − *F*_0_)/*F*_0_, where where F_n_ and F_0_ is the average intensity at n^th^ frame and at resting state, respectively.

### MEA preparation

MEA measurements were performed using an MEA 2,100-Lite Amplifier (Multi Channel Systems MCS GmbH). The 6-well MEA device was fabricated from a manufacturer (Multi Channel Systems MCS GmbH), and it contained 9 embedded 30◻μm diameter TiN electrodes per well with 200◻μm spacing between electrodes and six reference electrodes. The MEA device was coated with 0.1 mg/ml Poly-D-Lysine (Sigma) and Matrigel for cell culture preparation. The cell seeding density for cultures in MEAs was 0.4-0.6 M/cm^2^. Measurements were performed at a sampling rate of 10 kHz for 5 min at 37 °C with a sealed cover to keep CO_2_ concentration stable. Media was replenished every other day.

### MEA recording, and spike/burst detection

Neuronal activity was analyzed by Multi-Channel Analyzer software (Multi Channel Systems MCS GmbH), Python (3.9.7), and MATLAB. Raw data were filtered using a 2nd order Butterworth high pass filter with 200 Hz cutoff frequency. Action potentials were detected as spikes by a threshold of 4◻×◻standard deviations for both rising and falling edge from the noise magnitude distribution. Spikes were only detected by active electrodes which were defined by electrodes containing at least 5 spikes/min. The following criteria were used to define bursts. A burst must consist of at least 4 spikes, last at least 50 ms long, and must be separated from another burst by at least 100 ms. To start and end a burst, the interval between the first two and the last two spikes should be less than 50 ms, respectively.

### Synchrony index

The synchrony of spike trains between electrodes was assessed through cross-correlation for discrete functions (Pagan-Diaz et al., 2020). There are total nine electrodes and spike train data for each electrode was cross-correlated with every other electrode. The synchrony index was achieved by averaging of 36 possible combinations. The cross-correlation was proceeded with zero lag. The synchrony index, 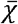, of nine electrodes is described as

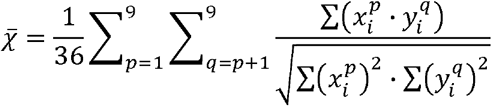

where *x* and *y* are spike trains consisting of 0 (no spike) and 1 (spike), *p* and *q* are the *p*^th^ and *q*^th^ electrode, and *i* is the *i*^th^ discrete time. 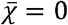 and 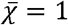 represent completely asynchronous and synchronous, respectively.

### Statistical analysis

SAS (9.4) and R (4.0.3) were used for statistical analysis. p◻<◻0.05 was considered statistically significant. Data were considered normally distributed when the absolute value of the skewness and kurtosis was less than 1 and 2, respectively. In the case of non-normal distribution, a power transform was used to transform data to meet the normality conditions. Actin intensity, muscle contraction amplitudes, calcium signal, and astrocyte number in response to the glia inhibitor were evaluated by two-sample t-test (RM vs. CM, control vs. BTS 10 μM, and control vs. glia inhibitor). Neuron number, synapse number, vesicle accumulation, and astrocyte number were evaluated by two-way ANOVA with day (day 2 or 3 vs. day 9) and treatment (RM vs. CM) as factors. The MEA outcomes of the BTS study were analyzed using repeated measures three-way ANOVA with cohort as a blocking variable, day (day 2 to 9) entered as a within-subjects, muscle treatment (2 levels: RM vs. CM) as a between-subjects factor, and drug (2 levels: control vs. BTS) entered as a between-subjects factor. Similarly, the MEA outcomes of the glia reduced study were analyzed using repeated measures three-way ANOVA with cohort as a blocking variable, day (day 2 to 9) entered as a within-subjects, muscle treatment (2 levels: RM vs. CM) as a between-subjects factor, and astrocyte composition (3 levels: presence vs. absence vs. absence with astrocyte releasate (Ast-RM and Ast-CM)) as a between-subjects factor. For the burst rate (BTS and glia reduced study) and synchrony index (glia reduced study), aligned rank transform was used for the non-parametric test since data were considered non-normally distributed. Post-hoc pairwise differences between means were performed using Fisher’s least significant difference test.

## Results

### Contracting muscle conditioned media enhances neuronal activity measured by microelectrode arrays

Consistent with our previous MEA study with C2C12 mouse myoblast cell line and mouse embryonic stem cell-derived neuronal culture (Aydin et al., 2020), CM from primary skeletal muscle cells increased spike and burst rates of primary hippocampal neurons across days (Fig 1A and 1B). The general pattern of development of spike trains over time in RM was consistent with other studies using primary hippocampal cells and primary sensory neurons at a similar cell seeding density (Biffi et al., 2013; Black et al., 2018). Significant differences in spike rates were observed between days (F_7, 80_ = 19.4, p < 0.001) and between RM versus CM treatments (F_1, 80_ = 202.2, p < 0.001). The interaction between day and treatment was also significant (F_7, 80_ = 23.7, p < 0.001). Similar to spike rate, burst rate also showed significant effects of day (F_7, 72_ = 10.5, p < 0.001), treatment (F_1, 72_ = 38.3, p < 0.001), and interaction between the two (F_7, 72_ = 21.1, p < 0.001). The interactions were caused by a different pattern of results for earlier time-points (days 2-7), as compared to later time-points (days 8 and 9). At the early time-points, CM had a higher spike and burst rates, but at later time-points the difference in them between RM and CM was reduced, absent, or reversed.

**Fig 1.**
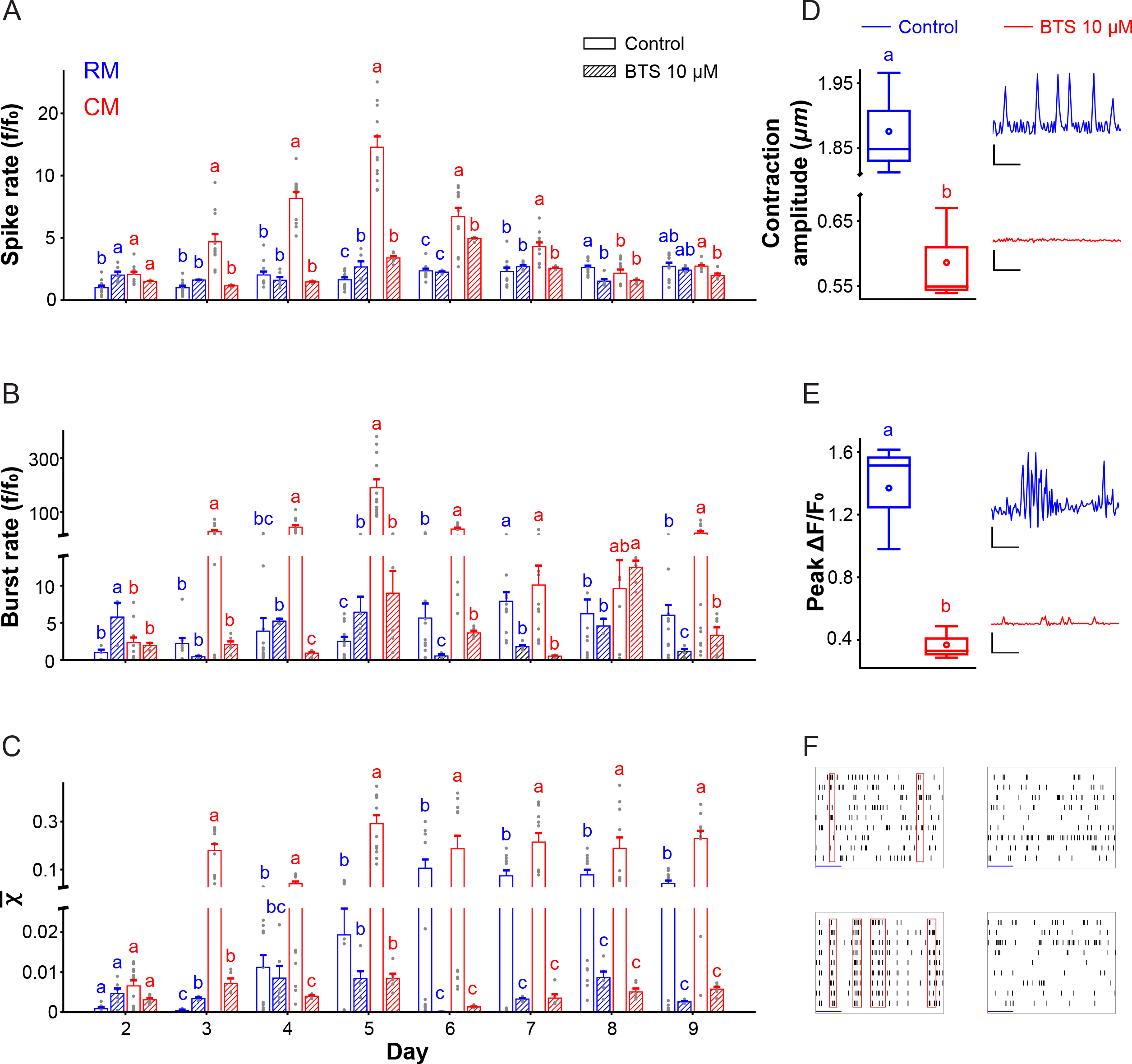
Muscle contraction conditioned media increases hippocampal neuronal activity. (A) Primary mouse hippocampal cells were cultured on a multi-electrode array (MEA) for 9 days with the following treatments: normal media (RM), muscle contraction conditioned media (CM), RM with N-benzyl-p-toluene sulphonamide (RM-BTS), CM with BTS. Average number of spikes per second (± SEM) across all electrodes normalized to average value for RM on day 2 are shown (n = 12 control, 6 BTS). Different lowercase letters indicate significant differences within each day (p < 0.05). (B) Average number of bursts per minute normalized to average value for RM on day 2. (C) Average synchrony index. (D) Average muscle contraction amplitudes between control and 10 μM BTS (left). Skeletal muscle contraction patterns in the presence of 0 (top right) and 10 μM BTS (bottom right). Scale bar in x, y: 1 s, 1 μm. (E) Peak fluorescent changes between control and 10 μM BTS. Traces of calcium dynamics in the presence of 0 (top right) and 10 μM BTS (bottom right) (n = 3). Scale bar in x, y: 2 s, 1 ΔF⁄F0. (F) Raster plots representing spike trains from nine electrodes from RM (top left), RM with BTS (top right), CM (bottom left), and CM with BTS (bottom right) on day 9. Each line and the red box represent a single firing, and synchronous burst, respectively. Scale bar: 2 s.

Similar to spike rate and burst rate, CM also caused neurons to fire more synchronously as compared to RM (Fig 1C). A two-way ANOVA showed a significant effect of day (F_7, 75_ = 175.7, p < 0.001), treatment (F_1, 75_ = 4083.4, p < 0.001) and the interaction between the two (F_7, 75_ = 231.1, p < 0.001). However, unlike spike rate and burst rate which showed greater differences between CM and RM at initial time-points than later time-points, the synchrony index showed the reverse pattern, with greater differences at the later time-points and no difference at the early time-points when little synchronous firing occurred.

Having shown that CM increases spike rate, burst rate and synchronous firing of primary hippocampal neurons in culture, we next wanted to evaluate whether contraction of the muscles was necessary for the MEA effects. The alternative is that muscle cells release neuro-active factors regardless of whether they are contracting. Hence, we repeated the experiment except we treated the muscle cells with a contraction inhibitor before collecting the media. We used a known skeletal muscle myosin II inhibitor, N-benzyl-p-toluene sulphonamide (BTS). BTS weakens myosin's interaction with F-actin and *ex vivo* studies found 10 μM of BTS suppresses force production by 60% (Cheung et al., 2002; Pinniger et al., 2005). Consistent with these results, the amplitudes of muscle contraction were reduced by 69% with BTS *in vitro* (t_4_= 20.7, p < 0.001; Fig 1D). Moreover, we performed calcium imaging of the skeletal muscles *in vitro* (see details in Materials and methods). Calcium dynamics in muscle cultures show a 73% reduction between control and 10 μM of BTS (t_4_= 4.9, p =0.0083; Fig 1E). Widefield (Video S1 and S2) and calcium imaging (Video S3 and S4) of skeletal muscles are available as supplementary materials.

BTS prevented CM from increasing spike and burst rate, but had no effect on baseline spike and burst rate in RM. This suggests muscle cell contractions are required for CM to increase spike and burst rate. For spike and burst rate, all factors in the repeated measures ANOVA were significant including day (F_7, 212_ = 50.2, p < 0.001; F_7, 187_ = 24.3, p < 0.001), muscle treatment (RM vs. CM) (F_1, 31_ = 194.0, p < 0.001; F_1, 21_ = 48.8, p < 0.001), BTS treatment (F_1, 31_ = 101.2, p < 0.001; F_1, 21_ = 52.8, p < 0.001) and all interactions (all p < 0.001). Post-hoc comparisons showed no difference between RM and RM with BTS (p = 0.4516; p = 0.1607), whereas CM produced greater spike and burst rates as compared to CM with BTS (p < 0.001, p < 0.001). Finally, CM with BTS was not different from RM (p = 0.1272, p = 0.9548) or RM with BTS (p = 0.4553; p = 0.1643).

While BTS specifically reduced spike and burst rate in CM with no effect on RM, we observed a different result for the synchrony index. BTS completely obliterated the synchrony index when added to either RM or CM treatments. This was supported by a significant effect of day, muscle treatment, BTS treatment and all interactions in the overall repeated measures ANOVA. Post-hoc tests showed that BTS-RM and BTS-CM displayed near zero synchrony across the entire period. The synchrony indices in BTS-RM and BTS-CM groups were significantly lower than RM (p < 0.001, p < 0.001) and CM without BTS (p < 0.001, p < 0.001) collapsed across days. Taken together, this suggests BTS has a direct effect on the cell culture preventing synchrony and hence we cannot draw strong conclusions about whether contraction of muscles is required for increasing the synchrony index without further experimentation.

### Contracting muscle conditioned media promotes synaptogenesis

Primary hippocampal cells plated in culture form synapses over a period of days. Increased number or strength of synapses could explain the increased synchrony index in CM compared to RM. To quantify synaptic development in response to CM versus RM, we performed immunocytochemistry to count the number of synapses using co-localization of pre- and post-synaptic markers (see details in Materials and methods) on days 2 and 9 for CM and RM (Dzyubenko et al., 2016).

The results showed that CM expedites synaptic development compared to RM. This was supported by a significant effect of day (F_1, 20_◻=◻19.6, p◻<◻0.001), no main effect of treatment (RM vs. CM) (F_1,20_◻=◻0.77, p◻=◻0.3896), but a significant interaction between day and treatment (F_1, 20_◻=◻8.5, p◻=◻0.0087; Fig. 2A). Post-hoc tests indicated a significant increase in the synapse number from day 2 to 9 in RM (p < 0.001), but not in CM (p = 0.2953). On day 2, the synapse number in CM was significantly higher by 44 % compared to RM (p = 0.0145), but on day 9, no differences between CM and RM were detected (p = 0.1669). The confocal images of post-synapses, synaptic vesicles, and colocalization are shown in supplementary materials (Fig S1).

**Fig 2.**
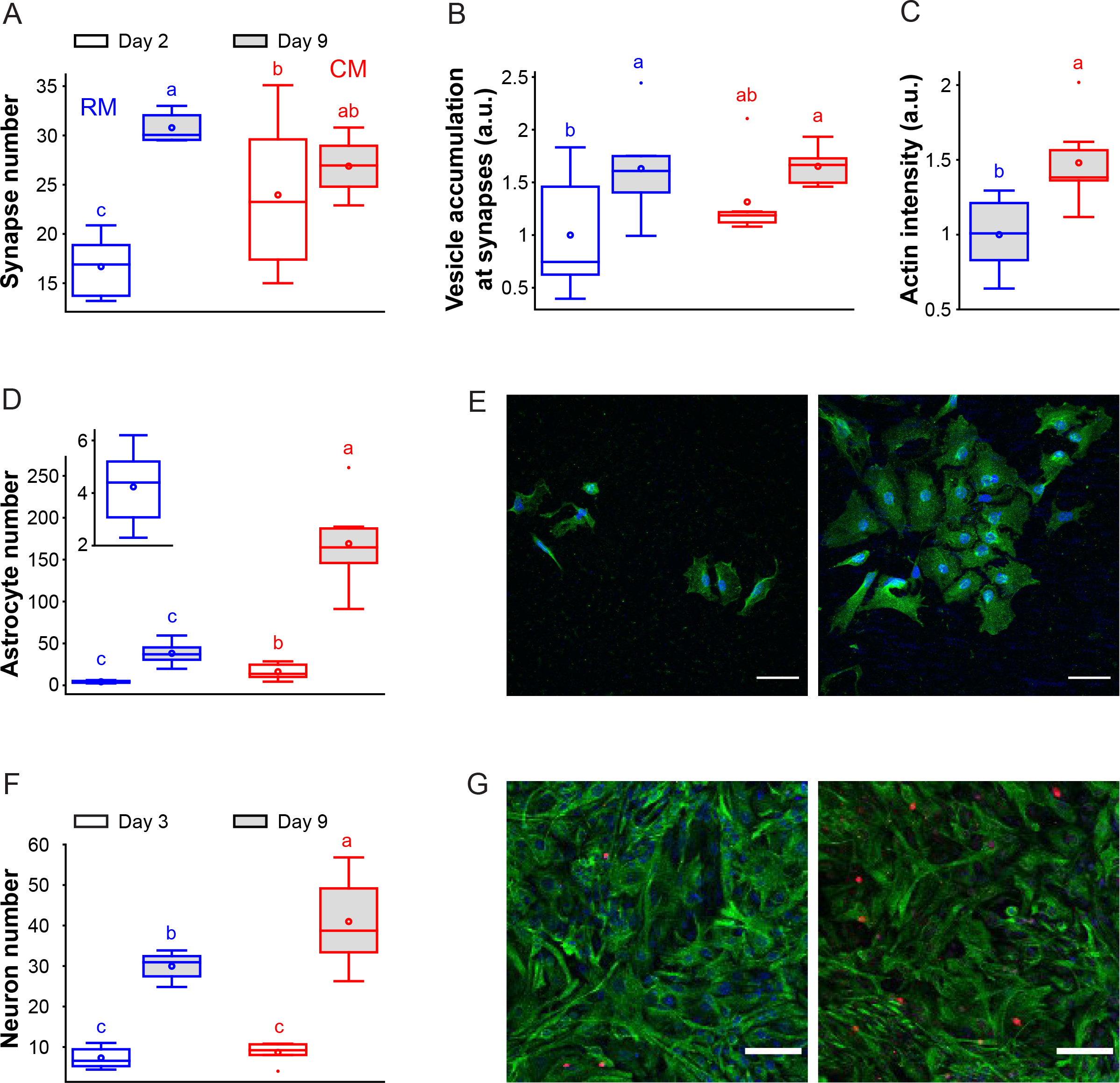
Muscle contraction conditioned media accelerates hippocampal neuron synapse development. (A) Primary mouse hippocampal cells were cultured for 9 days with normal media (RM) or muscle contraction conditioned media (CM). Average number of synapses (± SEM), as measured by immunohistochemical detection of pre and postsynaptic markers, are shown for RM and CM on days 2 and 9 (n = 6 per group). (B) Average vesicle accumulation per synapse on days 2 and 9. Data are shown normalized to average value for RM on day 2. (C) Average F-actin intensity on day 9 (F-actin levels were below limit of detection on day 2). Data are shown normalized to average value for RM. (D) Average number of astrocytes on days 2 and 9. The inset shows day 2 RM on a smaller scale. (E) Confocal images of astrocytes in the cultures on day 2 in RM (left) and CM (right) (S100β, green; DAPI, blue). Scale bar: 50 μm. (F) Average number of neurons on day 3 and 9 (n = 6 for day 3 and n = 8 for day 9). (G) Confocal images of neurons in the cultures on day 9 in RM (left) and CM (right) (NeuN, red; Nestin, green; DAPI, blue). Scale bar: 100 μm. Different lowercase letters indicate significant differences (p < 0.05).

### Contracting muscle conditioned media accrues filamentous actin at presynaptic terminals but does not significantly affect vesicle clustering

Functional synapses display an accumulation of vesicles and filamentous actin at the terminals (Fletcher et al., 1991). F-actin plays a critical role in clustering and transporting vesicles within the synapse (Pieribone et al., 1995; Kim and Lisman, 1999; Pechstein and Shupliakov, 2010; Peng et al., 2012). Hence, we wanted to determine whether CM affected vesicle accumulation and filamentous actin concentration at the synapse as a potential mechanism for the increased neuronal activity and synchrony observed in CM (Fig. 1A-C).

Following established methods (Dzyubenko et al., 2016) to quantify neurotransmitter vesicle clustering at the synapse, we measured average intensity of synaptophysin, a transmembrane protein for vesicles, co-localized with Bassoon, presynaptic nerve terminal marker. Results indicated vesicle clustering occurred at a similar rate in CM and RM. The ANOVA indicated a significant effect of day (F_1, 20_◻=◻7.6, p◻=◻0.0121), but no main effect of treatment (F_1, 20_◻=◻1.1, p◻=◻0.2990) or interaction (F_1, 20_◻=◻0.79, p = 0.3844; Fig 2B). On day 2, F-actin was not detected in either CM or RM so results are not shown. However, filamentous actin was detected on day 9 in both groups, and average F-actin intensity in CM was higher by 48% (t_10_= 2.9, p = 0.0152; Fig 2C).

### Muscle conditioned media from contracting muscles induces astrocyte proliferation

The numbers of astrocytes significantly increased by tenfold from day 2 to day 9 collapsed across RM and CM. CM consistently displayed fourfold greater numbers of astrocytes than RM on both days (Fig. 2D). This was reflected by a significant effect of day (F_1, 20_◻=◻145.0, p◻<◻0.001) and treatment (F_1, 20_◻=◻49.1, p◻<◻0.001) but no interaction (F_1, 20_ = 0.93, p = 0.3466). This suggests CM massively increases the proliferation of hippocampal astrocytes similar to the effect observed *in vivo* (Gardner et al., 2020).

### Muscle conditioned media from contracting muscles induces neuron proliferation

Number of neurons increased by 4.4-fold from day 3 to day 9 collapsed across RM and CM indicating significant neurogenesis in culture. No difference in the number of neurons were observed on day 3, but CM increased number of neurons by 40% as compared to RM on day 9 (Fig. 2F). This was reflected by a significant effect of day (F_1, 19_◻=◻180.5, p◻<◻0.001), treatment (F_1, 19_◻=◻8.6, p◻=◻0.0086), and interaction (F_1, 19_ = 5.5, p = 0.0295).

### Astrocytes regulate neuronal activity *in vitro*

To determine the role of astrocytes in the increased spike rate observed in hippocampal primary cultures exposed to CM versus RM, we repeated the MEA experiment in cultures with reduced astrocyte populations. To remove astrocytes from the primary hippocampal cell culture, we applied a glia inhibitor following established protocols (Liu et al., 2013) (see details in Materials and methods). Consistent with previous accounts, this resulted in an 81% reduction in the number of astrocytes in the culture (t_14_= 5.70, p < 0.001) (Fig 3A). Confocal images of astrocyte populations in control and astrocyte reduced culture are shown (Fig 3B).

**Fig 3.**
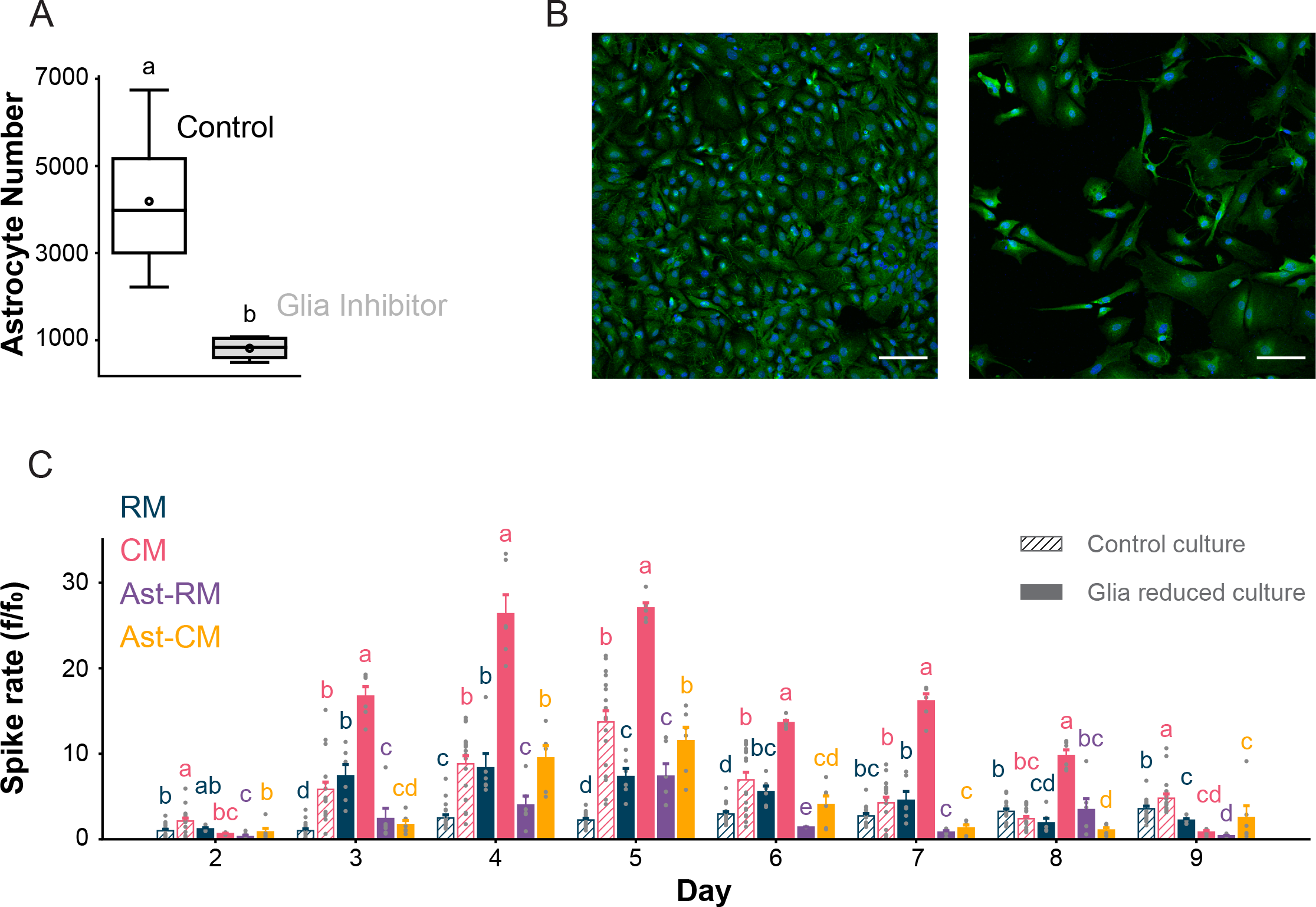
Astrocytes release factors that tame muscle-media induced neuronal activity. (A) Average number of astrocytes (± SEM) in control and glia reduced group on day 4 (n = 8 per group). (B) Confocal images of the control and glia reduced cultures on day 4 (S100β, green; DAPI, blue). Scale bar: 100 μm. (C) Average number of spikes per second normalized to RM on day 2 in the presence and absence of astrocytes (n = 18 for control and n=6 for glia reduced cultures). Different lowercase letters indicate significant differences within each day (p < 0.05). Results for burst rate and synchrony index are shown in supplementary materials (Fig S2 and Appendix S1).

MEA data were collected from normal hippocampal cultures in RM and CM as reference controls, as well as from cultures with reduced astrocytes in RM, CM, Ast-RM, and Ast-CM (see Materials and Methods section on “Collection of muscle and astrocyte media”). In the repeated measures analysis of mean spike rate, all main effects of muscle conditioned media (RM vs CM) (F_1, 53_ = 254.8, p < 0.001), presence or absence of astrocytes (F_2, 53_ = 170.6, p < 0.001), and day (F_7, 364_ = 131.7, p < 0.001) were significant, and all possible interactions between these factors were also significant (p < 0.001). The presence or absence of astrocytes factor includes 3 levels, presence, absence, and absence but with the releasate from astrocytes added back in (Ast-RM and Ast-CM groups; see statistical methods). Post-hoc tests indicated that CM increased spike rate relative to RM in normal hippocampal cultures consistent with the previous result (p < 0.001), but also in cultures with reduced astrocytes (p < 0.001). To our surprise, we found that a reduction of astrocytes increased spike rate in both RM and CM (p < 0.001), and the increase in CM with reduced astrocytes was an order of magnitude higher compared to the unaltered culture in CM (Fig 3C). Moreover, the increase in spike rate in CM relative to RM was greater for cultures with reduced astrocytes as compared to unaltered cultures as reflected by the significant interaction between muscle media and presence/absence of astrocytes (F_2, 53_ = 49.8, p < 0.001). Taken together, these results suggest astrocytes inhibit neuronal activity and CM increases their inhibitory function to counteract the excitatory effect of CM on neuronal activity.

To determine whether astrocytes mediate their inhibitory function through releasing factors into the media or whether they need to be physically present to exert their inhibitory effect, we included the Ast-RM and Ast-CM treatments. In these treatments, astrocytes were removed from the hippocampal culture but media from intact hippocampal cultures with astrocytes was added back in after being exposed to either RM or CM. RM and Ast-CM reduced spike rate relative to RM and CM when the hippocampal cultures were deprived of astrocytes. Moreover, the spike rate in the Ast-groups was similar to when the astrocytes are physically present in intact primary hippocampal cultures. This is supported by non-significant post-hoc test between groups where astrocytes were physically present versus absent but releasate added back in (p = 0.0537). Further, comparisons between normal cultures exposed to CM and astrocyte-deprived cultures exposed to Ast-CM showed no significant difference (p = 0.9597). Likewise, normal cultures exposed to RM showed no difference from astrocyte deprived cultures with Ast-RM (p = 0.68). These results suggest that astrocytes mediate their inhibitory effect through releasing factors into the media and do not need to be physically present in the culture to exert their influence.

## Discussion

Here we establish for the first time an *in vitro* platform to explore interactions between contracting primary muscle cells and primary hippocampal cells. One of the leading hypothesized mechanisms for pro-cognitive effects of exercise is that muscle contractions release factors that cross into the brain where they directly influence hippocampal cells involved in cognition (van Praag et al., 1999; Trejo et al., 2001; Wrann et al., 2013). This hypothesis is supported by our recent discovery that muscle contractions alone, through electrical stimulation of the sciatic nerve in anesthetized mice, are capable of increasing the generation of new astrocytes in the hippocampus and increasing the volume of the dentate gyrus (Gardner et al., 2020). The *in vitro* model developed herein adds to this literature by identifying a novel mechanism by which muscle cells may communicate with hippocampal cells. Muscle cells release factors that cause hippocampal neurons to become excited and hippocampal astrocytes to proliferate faster. The expanded astrocytes play a role in regulating neuronal excitability. Together this leads to a network that has overall greater excitability than in absence of the muscle signals, but also greater inhibition from astrocytes. The astrocytes thus tame the increased electrical activation of the circuit from the muscle factors in a way that leads to selective strengthening of coordinated activation patterns between neurons.

Astrocytes are well-known homeostatic regulators of neuronal activity. They directly modulate the ratio of excitatory and inhibitory synapses and neurotransmitter concentrations such as GABA and glutamate based on environmental needs. An *in vitro* study found the appearance of GABAergic and glutamatergic synapses by 24 hours when embryonic rat ventral spinal neurons were cultured on astrocytes as compared to 4 and 7 days, respectively in the neuron-only culture (Li et al., 1999). Furthermore, when astrocyte-conditioned media was supplemented in astrocyte-deprived situations, increases in GABAergic synapses, axon length (Hughes et al., 2010), and receptors (Diniz et al., 2014) were detected. Thus, astrocytes appear to release factors that increase GABAergic synapses and do not need to be physically adjacent to neurons to exert their inhibitory influence. This is consistent with their role in our study where media from neuronal cultures with astrocytes was capable of recapitulating the inhibitory effect of astrocytes in a neuronal culture without astrocytes physically present (Fig 3C). It is well established that astrocytes and astrocyte proliferation occur in response to epilepsy, and the evidence suggests astrocytes release factors such as gliotransmitters and tumor necrosis factor-alpha (TNF-α) that inhibit neuronal excitability and are protective against excitotoxicity (Verhoog et al., 2020).

A key finding was that muscle contractions are necessary for CM to influence spike rate and burst rate in the hippocampal cultures. When muscles were prevented from contracting by administering BTS, CM no longer produced the increased effects on spike and burst rate (Fig 1A and 1B). This adds important validity to the model since exercise involves mechanical forces and the hypothesis is that muscle contractions release factors that they otherwise would not release to communicate their status of engaging in physical activity to the hippocampus. We were able to make this conclusion because the effect of BTS on spike and burst rate was specific to CM, it had no impact when administered in RM, (i.e., BTS-RM spike and burst rate was similar to RM, but BTS-CM showed reduced spike and burst rate relative to CM). However, this was not true for the synchrony index where BTS appeared to directly eliminate synchronous firing of neurons whether in RM or CM (Fig 1C). Thus, we cannot be certain that the muscle contractions are necessary for the effect of CM on enhancing synchronous firing of action potentials. A method is needed that can prevent the muscle cells from contracting that does not directly interfere with any of the MEA outcomes.

In the context of whole organismal exercise, muscles communicate with hippocampal neurons while hippocampal neurons are involved in the sensorimotor processing in the brain that occurs during physical activity (Saraulli et al., 2017; Delezie and Handschin, 2018). Indeed, acute activation of the hippocampus is strongly correlated with running speed and repeated exercise training increases adult hippocampal neurogenesis and astrogliogenesis (van Praag et al., 1999; Redila and Christie, 2006; Clark et al., 2009; Erickson et al., 2011). Together with the *in vitro* data collected herein, the results suggest that muscle contractions contribute to the plasticity in the hippocampus by responding with signals that increase the number of new astrocytes to counterbalance the excitation that is likely intrinsic to the hippocampus involved in the sensorimotor response to physical activity.

Possibly related to the excitation of the hippocampus, whole organismal exercise produces a microenvironment in the hippocampus that is conducive for neurogenesis and synaptogenesis. Consistent with recent reports, results from the present study suggest that signals from muscle contractions likely contribute to the synaptogenic and neurogenic microenvironment. The fact that CM increased maturation of the hippocampal network, the formation of mature synapses and the formation of new neurons is consistent with such a role.

In this study, we used vesicle clustering and F-actin accumulation to quantify mature synapses. The presence of F-actin and vesicles at the terminal of presynaptic neurons is considered an indicator of a mature synapse. Previous *in vitro* studies have used the synaptic vesicle proteins, synapsin I and synaptophysin as markers to track the distribution of synaptic vesicles during the development of primary hippocampal neuron cultures. The synaptic vesicles change their distribution from being mostly in cell bodies to mostly at synapses as the hippocampal network matures. F-actin is known to be concentrated at presynaptic terminals to support the vesicles (Pieribone et al., 1995; Pechstein and Shupliakov, 2010; Peng et al., 2012) and to mediate their transportation (Kim and Lisman, 1999). Our observation that F-actin is concentrated more at presynaptic terminals in CM than RM implies that CM enhances the maturation of functional synapses with greater capacities to transport vesicles upon action potentials. This could explain why neuronal cultures plated on MEA exposed to CM displayed greater levels of synchronized firings of action potentials than RM because there were more mature synapses.

The increased number of astrocytes in the neuronal cultures exposed to CM may have contributed to the increased maturation of the hippocampal network by strengthening specific synapses and weakening others by pruning and inhibition. Mature synapses appeared earlier and were pruned earlier in cultures with more astrocytes as a consequence of exposure to CM. Whereas synapses continued to increase from day 2 to day 9 in RM, in CM they reached their peak around day 2 and were already in decline by day 9. Astrocyte-secreted proteins such as thrombospondins (Christopherson et al., 2005), hevin, and SPARC (Kucukdereli et al., 2011) may have promoted synaptogenesis. Astrocytes can directly eliminate synapses through MEGF10 and MERTK pathways which are two phagocytic receptors detecting signals from silent synapses (Chung et al., 2013, 2015).

Before conducting the present studies, we knew astrocytes increased in the hippocampus in response to exercise, and that muscle contraction alone was capable of recapitulating this effect. We observed the same phenomenon of increased astrocytes in the *in vitro* model, which justifies its use for exploring the role of increased astrocytes in the hippocampal response to muscle contractions. Because of the *in vitro* model, we now have a hypothesis for why astrocytes are responsive to muscle factors, they play an inhibitory role in taming the excitability of neurons that occurs in parallel with muscle contractions.

Future studies will build on the *in vitro* platform to explore potential reciprocal communication between muscle cells and hippocampal cells through a co-culture with shared media exchange. We are also interested in using the platform to explore the potential mechanism by which CM causes astrocytes to proliferate and hippocampal networks to mature faster. Finally, we are interested in identifying the bio-active factors released from the contracting muscles that influence the hippocampal cultures. In the future, such information could be used to reverse engineer treatments to recapitulate pro-cognitive effects of exercise in the absence of physical activity.

## Abbreviations

RM: Regular Media
CM: Contracting Muscle media
Ast-RM: Media taken from mixed astrocyte/neuron culture exposed to RM
Ast-CM: Media taken from mixed astrocyte/neuron culture exposed to CM
MEA: Microelectrode Array
BTS: N-benzyl-p-toluene sulphonamide

## Acknowledgements

We are grateful to Dr. Gelson Pagan-Diaz of the University of Texas for discussions of MEA, Jennie Gardner for general husbandry of animal subjects at the early stage of the study, Md Saddam Hossain Joy for discussions of the double-fluorescent label method, Carlos Renteria for discussions of calcium imaging and MATLAB code, and Meghan Connolly for discussions of neurogenesis. Lastly, we appreciate Dr. Onur Aydin of the University of Illinois for ideations and insightful discussions of the study.

## Figure captions

**Fig S1. Confocal images of synapses.** Post-synapses (PSD95) (top) and synaptic vesicle protein (synaptophysin) (bottom) in (A) RM and (B) CM on day 2. Orthogonal views of the colocalized channels in (C) RM and (D) CM. Synapses are indicated by the arrowhead. Scale bar: 50 μm.

**Fig S2. Regulation of neuronal activity by astrocytes on MEA.** (A) Burst rate normalized to RM on day 2. (B) Synchrony index in the presence and absence of astrocytes. Data are represented as mean ± SEM with data points (n = 18 (control), 6 (glia reduced culture)). Different lowercase letters indicate significant differences within each day (p < 0.05).

**Fig S3. Primary skeletal muscle contraction** *in vitro*. (A) Myotube with cross-striations indicated by the arrowhead (α-actinin, green; DAPI, blue). Scale bar: 10 μm. (B) Contracting myotube indicated by the arrowhead (left) on day 6. Scale bar: 100 μm. Contraction pattern of the corresponding muscle fiber (right). Scale bar in x, y: 1 s, 1 μm. (C) Lactate level between RM and CM measured by a colorimetric assay (n = 3 (RM), 6 (CM)). Different lowercase letters indicate significant differences (p < 0.05).

**Video S1. Primary skeletal muscle contraction in the absence of BTS.** Scale bar: 50 μm

**Video S2. Primary skeletal muscle contraction in the presence of 10 μM BTS.** Scale bar: 50 μm

**Video S3. Calcium dynamics of primary skeletal muscles in the absence of BTS.** Scale bar: 100 μm

**Video S4. Calcium dynamics of primary skeletal muscles in the presence of 10 μM BTS.** Scale bar: 100 μm

**Appendix S1. Supplementary experimental procedures and results**

