## Appendix S1 for "Astrocyte-mediated transduction of muscle fiber contractions synchronizes hippocampal neuronal network development"

### Appendix: supplementary experimental procedures and results

### Supplementary experimental procedures

#### L-lactate measurement

L-lactate concentration from primary skeletal myotubes was quantified using the Lactate Assay Kit (Sigma, MAK064), according to the manufacturer's instructions. The colorimetric method was chosen, and intensity was measured at 570 nm by a microplate reader (BioTek).

### Supplementary results

#### Astrocytes regulate burst rates and synchronous firings *in vitro*

In the repeated measures analysis of mean burst rate, all main effects of muscle conditioned media (RM vs CM) (F_1, 39_ = 466.0, p < 0.001), presence or absence of astrocytes (F_2, 39_ = 93.3, p < 0.001), and day (F_7, 330_ = 60.3, p < 0.001) were significant, and all possible interactions between these factors were also significant (p < 0.001). Likewise, significance differences were detected across muscle (F_1, 44_ = 76.6, p < 0.001), astrocyte (F_2, 44_ = 71.3, p < 0.001), day (F_7, 349_ = 39.4, p < 0.001) for the synchrony index. All possible interactions were significant (p < 0.001). Post-hoc tests indicated that CM increased both burst rate and synchrony index relative to RM in normal hippocampal cultures consistent with the previous result (p < 0.001), but also in cultures with reduced astrocytes (p < 0.001). Like the spike rate, a reduction of astrocytes increased both burst rate and synchrony index in both RM and CM (unaltered vs. glia reduced) (p < 0.001; p < 0.001). (Fig S2). Between astrocyte presence and releasate group, the Ast- group significantly decreased burst rate (p = 0.0349) and synchrony index (p = 0.0014) unlike the spike rate. This potentially suggests that unlike neuronal single firings, physical contact by astrocyte would be needed to accelerate neuronal circuit maturation such that action potentials occur in a burst form, and neurons communicate and fire synchronously.

#### Contracting muscle fibers release lactate into the media *in vitro*

Primary myoblasts were cultured from dissected hindlimb muscle tissue of 4-week-old mice and plated on flasks. Once the myoblasts differentiated into myotubes and myotubes started to contract, the media was changed, and the replenished media was collected every 24 hours. After collection, the media was supplemented with other reagents to be suitable for neuron culture (see details in Experimental procedures). This was used as the muscle conditioned media (CM). Regular media (RM), was the media used to replenish the myotube culture above (but never exposed to the myotubes), also with the addition of the same reagents. A confocal image of primary skeletal myotube with cross-striation in culture are shown (Fig S3A). In addition, a microscopic image of myotubes and corresponding muscle contraction pattern are shown (Fig S3B).

One of the components released by muscles during exercise *in vivo* is lactate. The concentration of the lactate proportionally changes depending on exercise intensity and duration (Goodwin et al., 2007; Menzies et al., 2010; Moraga et al., 2019). To confirm that contracting muscle cells release lactate *in vitro*, concentration of lactate was compared between RM and CM by a colorimetric assay. Based on the colorimetric test, the amount of lactate was approximately tenfold higher in CM than in RM (t_7_ = 33.8, p < 0.001) (Fig S3C). These results confirm that the contraction of myotubes, *in vitro*, release lactate into the media similar to how *in vivo* muscle contractions release lactate into the blood.
