## Supplementary figures and images for "Astrocyte-mediated transduction of muscle fiber contractions synchronizes hippocampal neuronal network development"

### Fig S1-S3

Fig S1

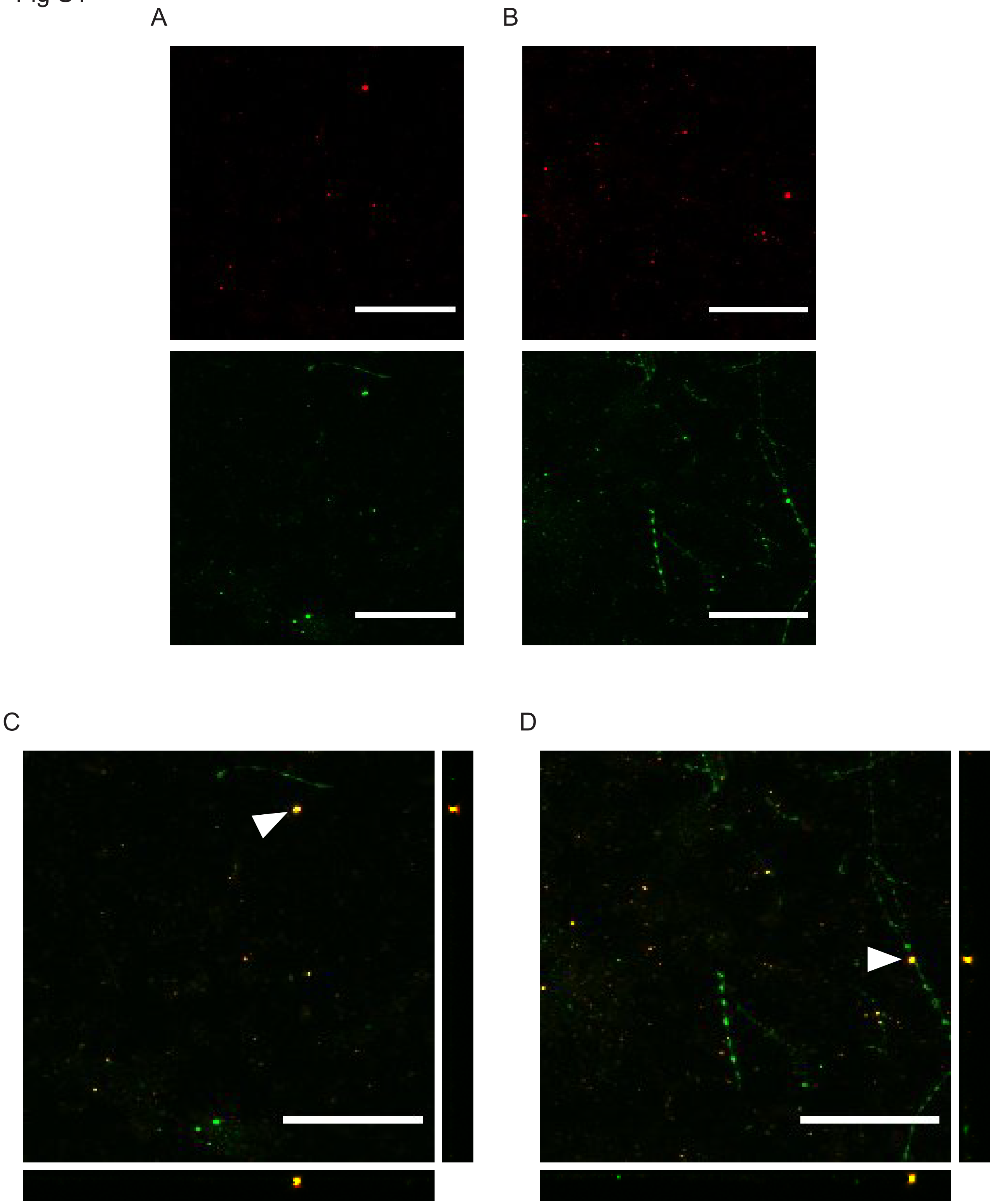

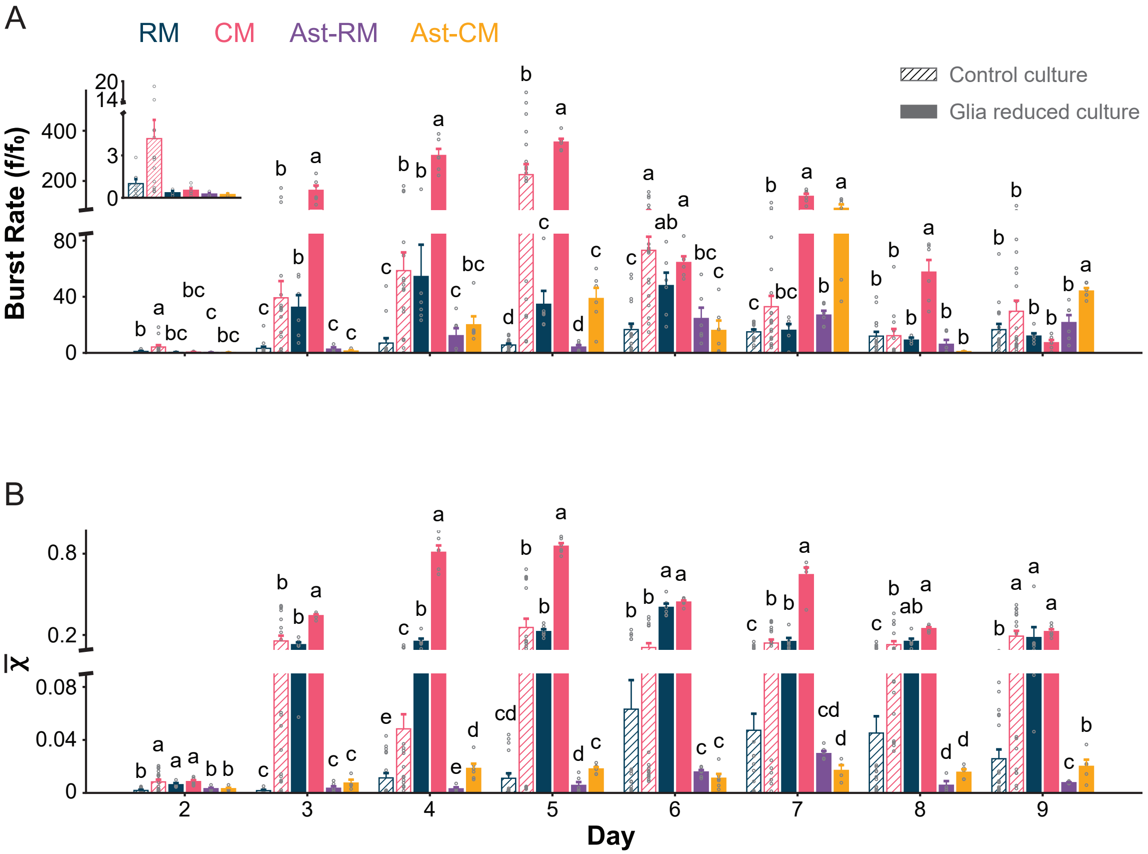

Fig S3

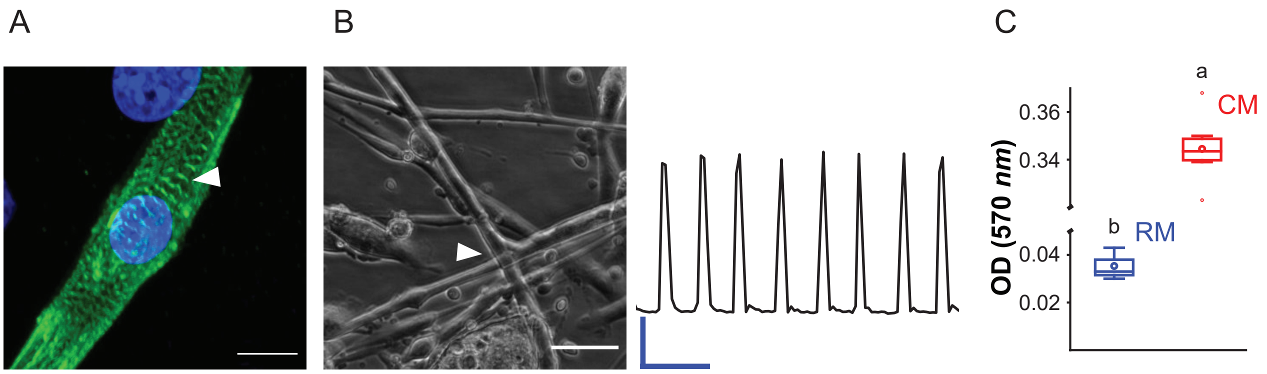
